# Estimating spatially and temporally complex range dynamics when detection is imperfect

**DOI:** 10.1101/586909

**Authors:** Clark S. Rushing, J. Andrew Royle, David J. Ziolkowski, Keith L. Pardieck

## Abstract

Species distributions are determined by the interaction of multiple biotic and abiotic factors, which produces complex spatial and temporal patterns of occurrence. As habitats and climate change due to anthropogenic activities, there is a need to develop species distribution models that can quantify these complex range dynamics. In this paper, we develop a dynamic occupancy model that uses a spatial generalized additive model to estimate non-linear spatial variation in occupancy not accounted for by environmental covariates. The model is flexible and can accommodate data from a range of sampling designs that provide information about both occupancy and detection probability. Output from the model can be used to create distribution maps and to estimate indices of temporal range dynamics. We demonstrate the utility of this approach by modeling long-term range dynamics of 10 eastern North American birds using data from the North American Breeding Bird Survey. We anticipate this framework will be particularly useful for modeling species’ distributions over large spatial scales and for quantifying range dynamics over long temporal scales.

## Introduction

The distribution of each species is determined by the interaction of multiple biotic and abiotic factors that vary across both space and time, including weather and climate (Barbet-Massin and Jetz 2014), habitat availability (Hill et al. 1999), physiological tolerances (Kearney and Porter 2009), and biotic interactions (Araújo and Luoto 2007). As a result, most species’ distributions are characterized by complex spatial and temporal patterns of occurrence, which combined with the large scales over which distributions change, present challenges for both the collection and analysis of data to quantify range dynamics (Elith et al. 2010). As habitats and climate change due to anthropogenic activities, there is an increasingly urgent need to develop species distribution models (SDMs) that can accurately and efficiently quantify complex range dynamics over large spatial and long temporal scales.

In response to this need, researchers have developed a range of SDM approaches that vary in their data requirements and analytical methods (Elith et al. 2010, Guillera-Arroita 2017). Although each of these methods has strengths and weaknesses, SDMs designed to quantify range dynamics require several key features. First, SDMs must include sufficient flexibility to quantify highly non-linear spatial patterns in occurrence probability. Although spatial variation in occurrence can in some cases be modeled using environmental covariates, residual spatial variation (which is likely common in most applications of SDMs at large spatial scales) can bias estimates of occurrence probability (Johnson et al. 2013, Guélat and Kéry 2018). Second, because occurrence probability at a given point in time is not independent of occurrence probability at earlier points in time, SDMs must explicitly account for temporal auto-correlation in occurrence probability (MacKenzie et al. 2003). If temporal auto-correlation is not accounted for within the SDM, occurrence dynamics are likely to appear more variable than they really are, leading to spurious conclusions about temporal variation range dynamics (Clement et al. 2016). Third, SDMs must uncouple true changes in occupancy from observation errors because the locations experiencing the largest changes in occupancy (e.g., range limits) are also the locations where errors arising from low detection probability are most likely (Tingley and Beissinger 2009, Guillera-Arroita 2017). Although progress has been made on accounting for each of these three issues (complex spatial variation, temporal auto-correlation, and imperfect detection) in SDMs, there are few modeling frameworks that address all three simultaneously.

Dynamic (or multi-season) occupancy models, which jointly estimate temporal change in the probability of occurrence and the probability of false-negative observations (MacKenzie et al. 2003), provide a natural framework that potentially meets each of above criteria (Kéry 2011). As a result, the use of occupancy models for species distribution modeling has grown in recent years (Guillera-Arroita 2017). At present, most occupancy-based SDMs have been implemented using one of several likelihood-based software programs, including programs PRESENCE (Hines 2006) and MARK (White and Burnham 1999) and the R package unmarked (Fiske and Chandler 2011). These programs allow users to fit several variations of the standard static or dynamic occupancy models (MacKenzie et al. 2003) using generalized linear models (GLMs) to estimate covariate effects on occupancy and detection. Although this GLM-based approach can account for imperfect detection and temporal auto-correlation in occupancy probability, these programs are restricted in their ability to model the complex non-linear patterns that typically characterize species’ distributions. In most cases, GLM-based models assume that spatial variation can be modeled as a linear or quadratic function of latitude and longitude, based on the assumption that species have a single center of high occupancy (Rich and Currie 2018). For many species, this assumption is not warranted and therefore GLM-based occupancy models are unlikely to provide accurate estimates of spatial variation in occupancy.

One alternative to the GLM approach is to estimate non-linear spatial variation in occupancy probability using generalized additive models (GAMs). An extension of GLMs, GAMs estimate the relationship between a response variable and a smoothed non-parametric function of covariates (Wood 2017). Because the shape of the smoothed functions is determined by the data rather than a parametric function, GAMs can estimate complex, nonlinear spatial patterns that are not accounted for by covariates (Hefley et al. 2017). Like GLMs, GAMs use link functions to accommodate response variables with normal or non-normal error distributions (e.g., binomial, Poisson), making it conceptually simple to extend occupancy models to estimate highly nonlinear effects of covariates (Kéry and Royle 2015). Although GAMs have been used to estimate species distributions in a number of contexts (e.g., Bled et al. 2013, Guélat and Kéry 2018), this approach has not been widely used in occupancy-based SDMs that account for both temporal dynamics and imperfect detection.

We developed a dynamic occupancy model that uses a spatial GAM to estimate non-linear spatial variation in occupancy not accounted for by environmental covariates. The model is flexible and can accommodate data from a range of sampling designs that provide information about both occupancy and detection. Output from the model can be used to create distribution maps and to estimate intuitive indices of range shifts (range center, range limits). We demonstrate the utility of this approach by modeling long-term (1972-2015) range dynamics of 10 eastern North American birds using data from the North American Breeding Bird Survey. For this application, we also use Gibbs variable selection (Dellaportas et al. 2002) to identify species-specific relationships between climate and occupancy. We anticipate this framework will be particularly useful for modeling species’ distributions over large spatial scales and for quantifying range dynamics over long temporal scales because of the improved fit to complex species distributions.

## Methods

### Model description

We assume that *j* = 1, 2,…*J* temporally or spatially replicated presence/absence surveys are conducted in *t* = 1, 2,…, *T* primary periods at *i* = 1, 2,…, *N* sampling locations. Further, we assume that the true (but latent) occupancy state of each site, denoted *z*_*i,t*_, is closed within each primary period but can change across primary periods. During each survey, the observed occupancy state of the focal species, denoted *h*_*i,j,t*_, is recorded (0 = species not observed, 1 = species observed). Our primary aim is to model temporal and spatial variation in the probability of occupancy *ψ*_*i,t*_ = *Pr*(*z*_*i,t*_ = 1) while accounting for imperfect detection. Below, we describe a Bayesian state-space formulation of this model that uses smoothing splines to model complex spatial and temporal auto-correlation in occupancy probability.

#### State model

In each primary period *t*, occupancy state is modeled as a Bernoulli random variable with probability *ψ*_*i,t*_:

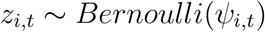

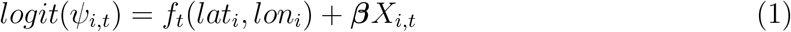

where *f*_*t*_(*lat*_*i*_, *lon*_*i*_) is a spatial smoothing function, ***β*** is a vector of slope coefficients, and ***X***_*i,t*_ is a matrix containing covariate values for route *i* in period *t*. The smooth function *f*_*t*_ is composed of basis functions *g*_*k*_ and their corresponding regressions coefficients *γ*_*k,t*_:

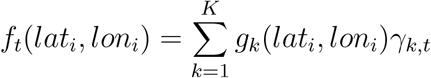

Different smooth functions can be chosen based on the structure of the data (Hefley et al. 2017, Wood 2017), providing a flexible and efficient means to model complex spatial variation in occupancy probability. The basis dimension *K* should be large enough to approximate the smooth function (i.e., avoid over-smoothing), though the exact choice of *K* is not critical because the degree of smoothing is determined primarily by a smoothing penalty term λ which penalizes against over-fitting (Wood 2017). In a Bayesian context, this penalization can be incorporated by specifying multivariate normal priors for the *γ*_*k*_ coefficients, with the precision matrix proportional to λ. Larger values of λ produce more constrained priors and thus more similar (i.e., more smooth) estimates of the *γ*_*k*_ coefficients (Wood 2017).

In combination with time-varying covariates ***X***_*i,t*_, allowing the basis function coefficients to vary across primary periods allows occupancy probability at each site to change over time. When *t* = 1, the *γ*_*k,1*_ coefficients are estimated using the Bayesian penalization approach described above. To account for temporal auto-correlation in occupancy, the basis function coefficients in periods *t* > 1 were modeled as temporally-correlated random effects:

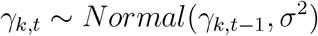

where *σ*^2^ is the variance among primary periods. Initial testing of our model indicated that the temporal random-effect formulation resulted in better mixing of the MCMC chains than the conventional dynamic occupancy model in which occupancy probability in years *t* > 1 is modeled as a function of extinction and colonization probabilities (MacKenzie et al. 2003). This formulation also accounts for temporal auto-correlation in occupancy probability without assuming that long-term dynamics are subject to equilibrium conditions.

#### Observation model

The observed status of each site during each survey (*h*_*i,j,t*_) is modeled as a function of both the latent state process (*z*_*i,t*_) and detection probability *p*_*t*_:

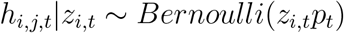

Covariates thought to influence the detection process (observer bias, weather, etc.) can be incorporated into this structure using a logistic link function on *p*_*t*_ (see below).

### Distribution maps and indices of range dynamics

When latitude, longitude, and annual covariate values are available at unsampled locations within a species’ range, posterior distributions of the predicted occupancy probability at those locations can be easily estimated from the posterior samples of the fitted model (Kéry 2011, Clement et al. 2016). Posterior estimates of range-wide occupancy probability can be used to visualize changes in species’ distributions and to quantify indices of range dynamics (Clement et al. 2016). Although many such indices are possible, we describe four that may be particularly relevant to quantifying range shifts. First, the mean occupancy probability of all map cells provides an index of changes in the proportion of area occupied (Clement et al. 2016). Second, the mean breeding latitude, estimated as the sum of the cell latitudes weighted by their occupancy probabilities and divided by the total occupancy probability across all cells, can be used to quantify shifts in the center of species range (Clement et al. 2016). Finally, annual indices of the northern/southern range limits can be estimated by sorting the map cells by latitude and then using a smoothing spline function to predict the latitude below/above which 99.9% of the total occupancy probability is located. Although not an absolute measure of the northern- and southern-most latitudes at which a species was found, this index provides a time series of relative change in the northern and southern range limits, which can be used to determine whether distributions have expanded or contracted over time. We provide code in Appendix S1 for estimation of these indices.

### Application

#### Data

We demonstrate the utility of this model by quantifying range dynamics of 10 eastern North American bird species: Red-bellied Woodpecker (*Melanerpes carolinus*), Fish Crow (*Corvus ossifragus*), Carolina Chickadee (*Poecile carolinensis*), Carolina Wren (*Thryothorus ludovicianus*), Blue-gray Gnatcatcher (*Polioptila caerulea*), Wood Thrush (*Hylocichla mustelina*), Golden-winged Warbler (*Vermivora chrysoptera*), Swainson’s Warbler (*Limnothlypis swainsonii*), Louisiana Waterthrush (*Parkesia motacilla*), and Kentucky Warbler (*Geothlypis formosa*). We selected these species because they exhibit a wide range of spatial and temporal complexity in range dynamics. Data for this analysis came from the North American Breeding Bird Survey (BBS), a large-scale citizen science program consisting of over 5500 roadside survey routes of which approximately 3000 are surveyed each May or June by trained volunteers (Sauer et al. 2015). The BBS was initiated in 1966, though only a small number of routes were surveyed during the early years. For this reason, we chose to use BBS data collected from 1972 to 2015 (Pardieck et al. 2016). Trained observers conduct 3-minute point counts at 50 regularly spaced stops along each 39.4km-long route. See Sauer et al. (2015) for more details regarding the BBS survey protocol. Prior to analysis, we converted the raw counts to stop-level presence/absence data.

To model spatial/climate relationships across the edge of each species’ occupied range, we subset all BBS routes with at least one detection of the focal species over the study period (i.e., routes occupied in at least one year). Next, we created a 2°-buffered convex hull around the occupied routes and included all routes within the buffered region.

Climate data was obtained from the University of East Anglia Climate Research Unit (CRU; Harris et al. (2014)). The CRU data contains global estimates of monthly surface climate variables for 0.5° grid cells (Harris et al. 2014). Following Clement et al. (2016), we converted the monthly temperature and precipitation estimates from the CRU data set into five ‘bioclim’ variables that have low correlation and are effective for modeling species ranges (Barbet-Massin and Jetz 2014). Specifically, for each grid cell we calculated the mean temperature, mean diurnal temperature range, mean temperature of the wettest quarter, annual precipitation, and precipitation of the warmest quarter for the 12 months preceding each BBS survey (i.e., June-May). All estimates were obtained using the ‘biovars’ function in the ‘dismo’ package (Hijmans et al. 2016) in program R (R Core Team 2016). Prior to analysis, each variable was scaled to mean = 0 and standard deviation = 1 and we extracted the annual climate values for the grid cell containing the first stop of each BBS route.

Because BBS routes are surveyed a single time each year, conventional methods for estimating detection probability from temporally replicated surveys are not possible. Instead, we adapted an occupancy model that uses the correlation between adjacent spatial replicates to estimate detection probability (Hines et al. 2010, 2014). In this model, occurrence is estimated at two scales: 1) the route-level (i.e., *z*_*i,t*_), and 2) the stop-level (denoted *y*_*j,i,t*_|*z*_*i,t*_). Hereafter we follow Clement et al. (2016) and refer to presence at the route-level as “occupancy” and presence at the stop-level as “availability”. Digital records of the raw 50-stop BBS data are only available from 1997-present. However, 10-stop summaries (sum of counts from stops 1-10, 11-20, 21-30, 31-40, and 41-50) are available for the entire BBS period (Pardieck et al. 2016). Initial testing indicated that estimates of the route-level occupancy did not differ when the model was fit using the full 50-stop data or the 10-stop summaries. Therefore, we chose to use the 10-stop data so that inferences could be made over the entire BBS time series.

#### State model

Spatial and annual variation in route-level occupancy probability was modeled using a modified version of Eq. 1:

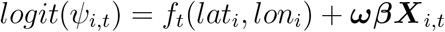

where ***ω*** is a vector of binary indicator variables determining whether each climate predictor is included in the model and ***X***_*i,t*_ is a matrix containing the annual climate values in year *t* at route *i*. Estimation of ***ω*** is described below. To capture non-linear relationships between climate and occupancy, the matrix ***X*** contained both the linear and quadratic terms for each of the 5 climate covariates. For the spatial smooth, we used a thin-plate regression spline of the latitude and longitude of each BBS route. For all species, we chose *k* = 60, which initial tests indicated was large enough to approximate the smooth function for all species considered.

At the stop level, availability is modeled as a first-order Markov process with parameters:

- *θ*_*i,j,t*_ = Pr(availability at stop *j*|route *i* occupied and stop *j* − 1 unavailable)
- 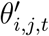 = Pr(availability at stop *j*|route *i* occupied and stop *j* − 1 available)

As noted by Hines et al. (2010), the first stop on each BBS routes has no predecessor and thus availability at stop 1 cannot be modeled using *θ* or *θ*′. Instead, we directly estimated the probability *π*_*t*_ that the first stop is available in year *t*:

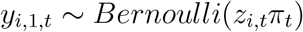

For the remaining stops, availability was modeled as:

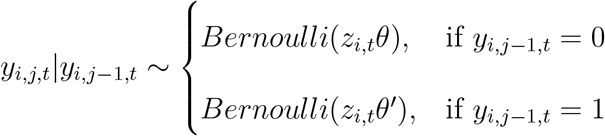

#### Observation model

Numerous factors could influence detection probability in BBS surveys. Observer experience and variation among observers are both known to influence detection of many species (Link and Sauer 2002). Weather conditions, particularly wind speed, may also influence detectability. For our analysis, we included wind speed scores recorded at the start of each BBS count using the Beaufort wind scale (Sauer et al. 2015). Between 2009 and 2015, a small number of BBS routes (n = 106) were surveyed using a modified protocol (RPID = 501) that incorporated time and distance information (Twedt 2015). Compared to the standard BBS protocol (RPID = 101), this time-distance protocol resulted in on average 10% fewer observations per survey (Sauer et al. *in prep*). To account for these effects, we modeled detection conditional on stop-level availability and route-level occupancy as:

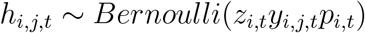

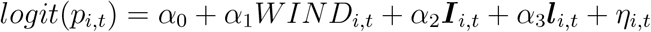

where *α*_0_ is an intercept term, *WIND*_*i,t*_ is the wind score, ***I***_*i,t*_ is a binary dummy variable indicating whether year *t* was an observer’s first year of service, ***l***_*i,t*_ is a dummy variable indicating the survey protocol used (0 = standard BBS survey, 1 = time-distance protocol), and *η*_*i,t*_ is a random observer effect.

#### Model selection

Given the large number of climate predictors in our model and the lack of *a priori* hypotheses about which predictors should influence the distribution of each species, each climate variable *m* was multiplied by a binary latent variable *ω*_*m*_ which determined whether the variable was included in the linear predictor. The posterior probability *Pr*(*ω*_*m*_ = 1) is then a measure of the relative importance of variable *m* (Kuo and Mallick 1998, Dellaportas et al. 2002, Ntzoufras 2002). For the linear effect of each climate variable, we assumed mutually independent Bernoulli priors:

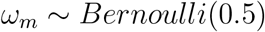

For the quadratic terms, we enforced marginality by setting:

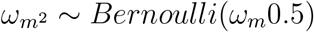

where *m*^2^ is the quadratic term associated with the linear term *m*. Thus, quadratic terms could only enter the model if the corresponding linear term is also in the model. To ensure good mixing of the *ω*_*m*_ parameters, we used Gibbs Variable Selection (GVS; Dellaportas et al. 2002) to create joint prior distributions for the *β*_*m*_ parameters conditional on *ω*_*m*_:

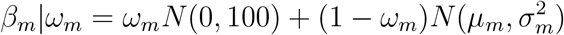

where *N* (0,100) is a non-informative normal prior when *ω*_*m*_ = 1 and 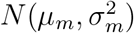 is a pseudo-prior sampled when *ω*_*m*_ = 0. We estimated *µ*_*m*_ and 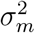 by running the correlated detection model in PRESENCE (Hines 2006) using the first 10 years of BBS data for each species and including linear and quadratic effects for all five climate predictors.

#### Modeling fitting and indices of range shifts

We fit the models in JAGS (Plummer 2012) called from R using the jagsUI package (Kellner 2015). As described above, we specified multivariate normal priors for the GAM smooth coefficients, Bernoulli priors for the indicator variables, and vague normal priors for the *β* coefficients. For all other parameters, we specified appropriate vague priors. See Data S1 for model code and specification details. We assessed goodness-of-fit using posterior predictive checks (PPC). Because conventional PPC metrics are inappropriate for binary occupancy data (Kéry and Royle 2015), we used each posterior estimate from the fitted model to simulate the expected number of routes with each of the 32 possible detection histories in each year. We used a Freeman-Tukey statistic (Brooks et al. 2000) to measure the discrepancy between the observed/simulated and predicted detection history frequencies and we report the Bayesian P-value from these tests.

For each species, we created annual distribution maps and range shift indices using the climate covariate values, latitude, and longitude for each 0.5° raster cell within the same buffered convex hull used to subset BBS routes. Posterior distributions of the predicted annual occupancy probability in each cell were estimated using the posterior samples for each model parameter. From these distributions we then estimated posterior distributions for the four indices described above (proportion of area occupied, mean breeding latitude, and northern/southern range limits) and report the mean and 95% credible interval for each.

## Results

By allowing for complex variation in occupancy while simultaneously penalizing against over-fitting, the spatial GAM was able to model both highly complex and relatively simple distributions. For example, species like Fish Crow and Swainson’s Warbler have complex spatial distributions that do not exhibit simple linear or quadratic relationships with latitude and/or longitude. Occupancy probabilities for the Fish Crow were high along the Atlantic and Gulf coasts as well as in the southern Mississippi River valley, resulting in a U-shaped distribution with no clear range center (Fig. 1A). The distribution of Swainson’s Warbler was also complex, with three distinct areas of high occupancy and low occupancy in between (Fig. 1B). In contrast, the distributions of some species, including Red-bellied Woodpecker and Carolina Chickadee, were less complex, with a large central area of high occupancy with declining occupancy along the periphery of the range (Fig. 1C-D).

**Figure 1:**
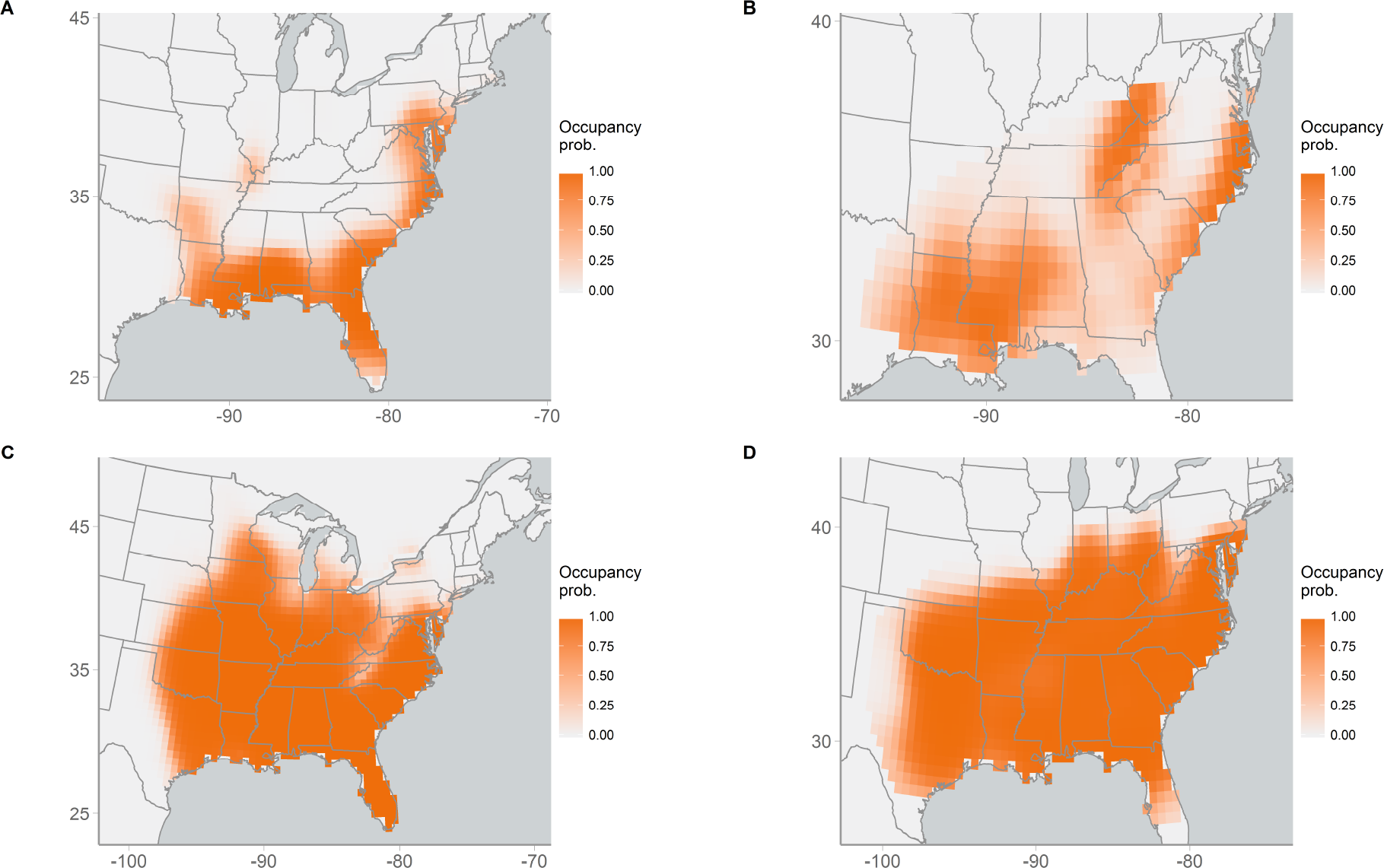
Distributions of (A) Fish Crow, (B) Swainson’s Warbler, (C) Carolina Chickadee, and (D) Eastern Towhee. Maps show the predicted probability of occupancy in 1972 for each species.

The model was also able to quantify temporal changes in occupancy probability over the 43 year time period of the BBS data, revealing similarities and differences in regional dynamics in several species. For example, Louisiana Waterthrush, Kentucky Warblers, Wood Thrush, and Golden-winged Warblers all experienced large declines in occupancy probability in the eastern portions of their range, especially in the northeastern United States and Appalachia, but were relatively stable or increasing in the mid-western United States (Fig. 2). These species differed, however, in occupancy trends in the southeastern United States, with Louisiana Waterthrush and Kentucky Warblers showing modest increases in occupancy probability and Wood Thrush and Golden-winged Warblers declining in occupancy probability.

**Figure 2:**
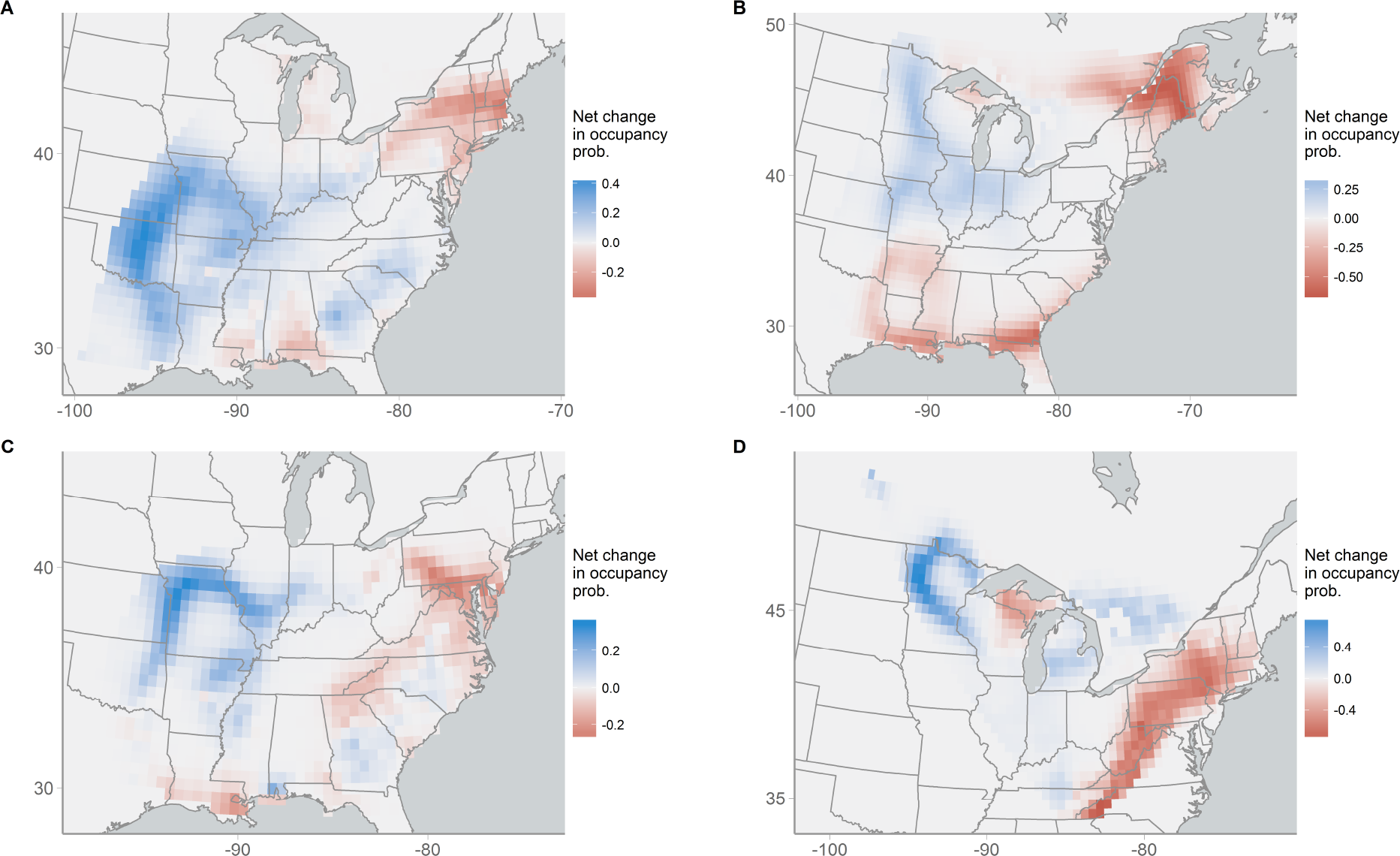
Net change in occupancy probability (*Ψ*_2015_ − *Ψ*_1972_) for (A) Louisiana Waterthrush, (B) Brown Thrasher, (C) Kentucky Warbler, and (D) Golden-winged Warbler. Blue indicates areas that have increased in occupancy probability; red indicates areas that have decreased in occupancy probability.

Indices of range shifts from our model indicate that some species have undergone distributional shifts over the past four decades. For example, the northern range limit of Blue-gray Gnatcatchers shifted northward by ~1.2° latitude and the mean breeding latitude shifted northward by ~1° (Fig. 3A-B). Interestingly, this species has shown small (~0.2°) southward expansion at the southern edge of its range and as a result, the proportion of area occupied has increased over time (Fig. 3C). The indices were also able to capture transient dynamics in distributional shifts. Northern populations of Carolina Wren, for example, experienced large declines in occupancy probability in the late 1970’s, resulting in a contraction of the northern range limit and mean breeding latitude by ~1° latitude (Fig. 3E-F). This contraction was temporary, however, with these populations subsequently experiencing a sustained northward expansion extending ~1.7° beyond their initial northern range limit (Fig. 3D).

**Figure 3:**
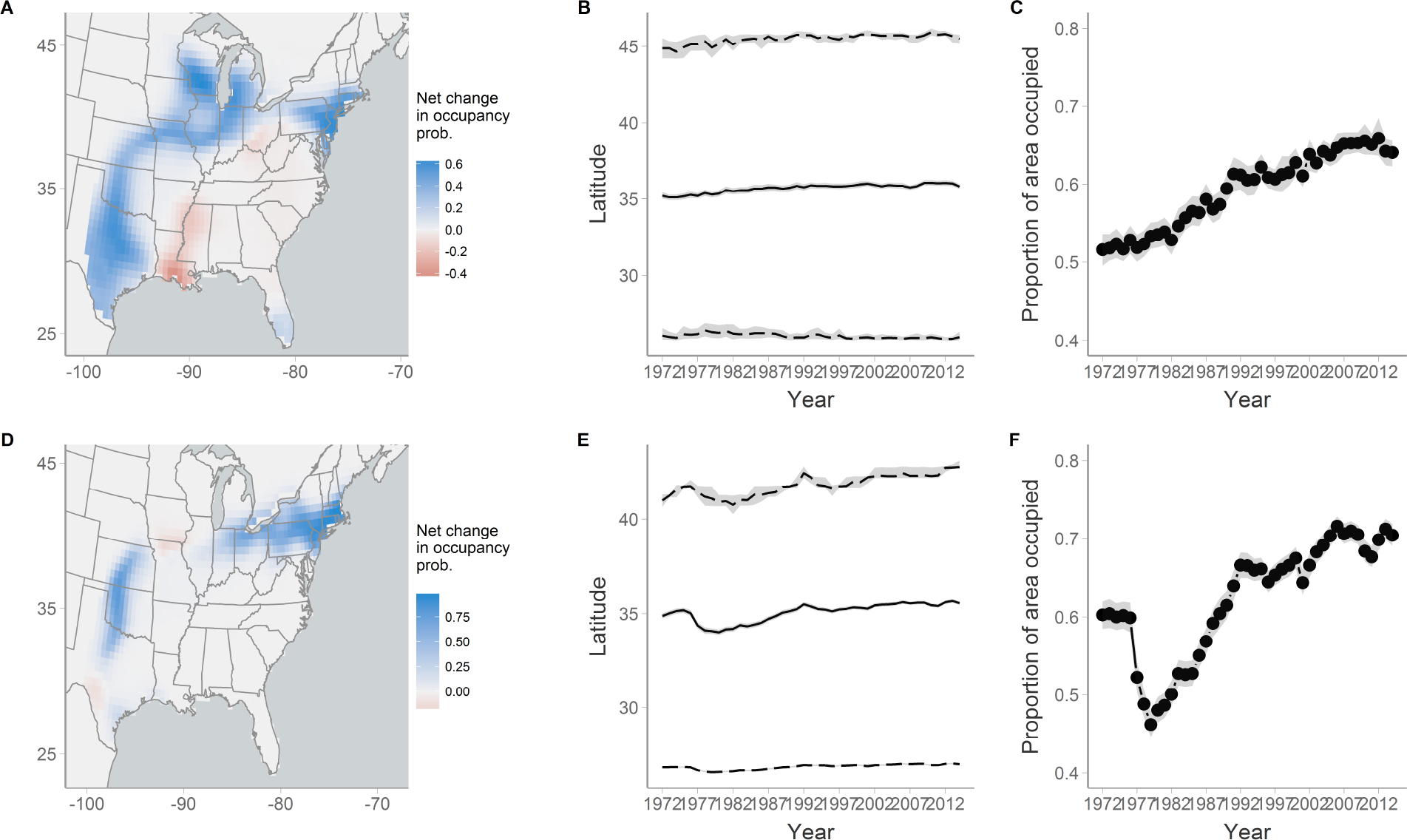
Net change in occupancy probability, latitudinal range indices, and proportion of area occupied for (A-C) Blue-gray Gnatcatcher and (D-F) Carolina Wren. For latitudinal range indices, solid lines indicate the mean breeding latitude and dashed lines indicate the northern and southern range limit. Gray ribbons represent the 95% credible intervals.

## Discussion

The distributions of most species are characterized by complex and dynamic variation in occurrence. Species distribution modeling seeks to relate this variation to environmental covariates and extrapolate these relationships to unsampled sites and times. Because habitats and species-habitat relationships change across both space and time, conventional GLM-based models rarely capture the inherent complexity of species distributions, especially when inferences are made across large spatial or long temporal scales. Here, we demonstrate a novel occupancy-based SDM that combines environmental predictors with a spatial GAM to model covariate relationships and complex, non-linear spatial variation in occupancy probability while accounting for imperfect detection.

Application of this model to 10 North American bird species demonstrates the utility and flexibility of our approach for species’ distribution modeling. Unlike parametric models that use low-order polynomials to capture spatial variation in occupancy, the spatial GAM derives non-parametric occupancy patterns during estimation while avoiding overfitting through the incorporation of a smoothing penalty term. This balancing of complexity and smoothing ensures that the model allows for, but does not impose, complex spatial variation in occupancy. This formulation provides an efficient and flexible method to model large-scale spatial variation in occupancy probability, for example allowing us to model both the complex distributions of Fish Crow and Swainson’s Warbler and the relativity more simple distributions of Carolina Chickadees and Red-bellied Woodpeckers using a common model structure.

Modeling the GAM coefficients as temporally-correlated random effects also allowed us to explicitly model changes in occupancy probability over time. In our analysis of BBS data, the model uncovered interesting similarities and differences in the range dynamics of several species that inhabit forest habitats in the eastern United States, including Louisiana Waterthrush, Wood Thrush, Kentucky Warblers, and Golden-winged Warblers. In the case of Wood Thrush and Golden-winged Warblers, this regional variation in occupancy dynamics is consistent with differences in demographic rates among the regions (Rosenberg et al. 2016, Rushing et al. 2016), suggesting that our occupancy model was able to capture spatial variation in population dynamics.

The indices of latitudinal range dynamics from our model also documented range expansions of Blue-gray Gnatcatchers and Carolina Wrens, demonstrating the utility of these metrics for quantifying range shifts over long temporal scales. The ability to document shifts at range margins while accounting for imperfect detection is particularly important given that these locations are likely to experience the largest changes in occupancy but also have the lowest detection probability. Application of this framework to a larger pool of species could indicate whether range shifts provide a consistent fingerprint of habitat or climate change. It may also be possible to quantify range shifts of entire groups of species by creating composite versions of our indices, which would be particularly useful for testing hypotheses about which traits promote or impede the ability of species to respond to habitat and climate change.

Our model differs from the conventional dynamic occupancy model in that we did not directly estimate change in occupancy as the result extinction/colonization processes. Under the extinction/colonization formulation, occupancy probability at a given location *i* will converge on the stable-state occupancy distribution defined by 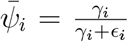, where *γ*_*i*_ and *ϵ*_*i*_ are the colonization and extinction rates at location *i* (MacKenzie et al. 2017). Initial testing of our model using the extinction/colonization formulation indicated that extinction rates were very low relative to colonization rates for most species, resulting in non-negligible occupancy probabilities (> ~8-10%) in the final year of our analysis at locations where the species were never detected. We suspect this may be a common issue when fitting dynamic occupancy models at large spatial and long temporal scales. Because our formulation does not make any equilibrium assumptions, it may be a better approach when the goal is to document range shifts. In some cases, using the conventional dynamic occupancy model with spatial variation in extinction/colonization probabilities may be preferable, particularly when the focus is on mechanistic understanding of range dynamics. Additionally, GAMs generally perform poorly when predicting outside of the data used to fit the model so the conventional occupancy model may be more suitable when the goal is to predict future distributions.

The model presented in this paper builds off of recent advances in the development of occupancy-based SDMs that account for both imperfect detection and temporal auto-correlation in occupancy (Clement et al. 2016). Our model extends this framework to account for complex spatial and temporal variation in occupancy probability through the use of a hierarchical Bayesian model with a spatial GAM. Accounting for complex spatial structure in SDMs is an active area of research (Reich et al. 2006) and other methods exist for handling spatial structure in occupancy models, particularly through the use of conditional autoregressive (CAR) modeling (e.g., Bled et al. 2011, Johnson et al. 2013, Guélat and Kéry 2018). Both CAR and GAM approaches have been shown to reduce bias in SDM models (Guélat and Kéry 2018), though the GAM approach is generally more computationally efficient, produces more precise parameter estimates, and can be fit using most popular Bayesian software programs, including JAGS, WinBUGS (Lunn et al. 2000), NIMBLE (Valpine et al. 2017), and STAN (Carpenter et al. 2017). These two approaches are not mutually exclusive though and future work integrating GAM and CAR models in occupancy-based SDMs is likely to improve inferences about past and future range dynamics.

## Acknowledgements

The authors thank J. Hines, M. Clement, J. Sauer, and N. Hostetter for assistance with model development and M. Kery for comments that improved earlier drafts of this manuscript. We also thank the thousands of BBS volunteers for collecting the data that made this project possible.

## Author contributions

CSR, JAR, and KLP concieved of the study with input from DJZ. CSR and JAR developed the methods. CSR carried out all analyses and drafted the manuscript. All authors contributed to subsequent revisions.

## Competing interests

The authors declare no competing interests.

## References

Araújo, M. B., and M. Luoto. 2007. The importance of biotic interactions for modelling species distributions under climate change. Global Ecology and Biogeography 16:743–753.

Barbet-Massin, M., and W. Jetz. 2014. A 40-year, continent-wide, multispecies assessment of relevant climate predictors for species distribution modelling. Diversity and Distributions 20:1285–1295.

Bled, F., J. D. Nichols, and R. Altwegg. 2013. Dynamic occupancy models for analyzing species’ range dynamics across large geographic scales. Ecology and Evolution 3:4896–4909.

Bled, F., J. A. Royle, and E. Cam. 2011. Hierarchical modeling of an invasive spread: The Eurasian Collared-Dove *streptopelia decaocto* in the United States. Ecological Applications 21:290–302.

Brooks, S. P., E. A. Catchpole, and B. J. Morgan. 2000. Bayesian animal survival estimation. Statistical Science:357–376.

Carpenter, B., A. Gelman, M. D. Hoffman, D. Lee, B. Goodrich, M. Betancourt, M. Brubaker, J. Guo, P. Li, and A. Riddell. 2017. Stan: A probabilistic programming language. Journal of Statistical Software 76.

Clement, M. J., J. E. Hines, J. D. Nichols, K. L. Pardieck, and D. J. Ziolkowski. 2016. Estimating indices of range shifts in birds using dynamic models when detection is imperfect. Global Change Biology 22:3273–3285.

Dellaportas, P., J. J. Forster, and I. Ntzoufras. 2002. On Bayesian model and variable selection using MCMC. Statistics and Computing 12:27–36.

Elith, J., M. Kearney, and S. Phillips. 2010. The art of modelling range-shifting species. Methods in Ecology and Evolution 1:330–342.

Fiske, I., and R. Chandler. 2011. Ummarked: An R package for fitting hierarchical models of wildlife occurrence and abundance. Journal of Statistical Software 43:1–23.

Guélat, J., and M. Kéry. 2018. Effects of spatial autocorrelation and imperfect detection on species distribution models. Methods in Ecology and Evolution:n/a–n/a.

Guillera-Arroita, G. 2017. Modelling of species distributions, range dynamics and communities under imperfect detection: Advances, challenges and opportunities. Ecography 40.

Harris, I., P. Jones, T. Osborn, and D. Lister. 2014. Updated high-resolution grids of monthly climatic observations–the CRU TS3. 10 Dataset. International Journal of Climatology 34:623–642.

Hefley, T. J., K. M. Broms, B. M. Brost, F. E. Buderman, S. L. Kay, H. R. Scharf, J. R. Tipton, P. J. Williams, and M. B. Hooten. 2017. The basis function approach for modeling autocorrelation in ecological data. Ecology 98:632–646.

Hijmans, R., S. Phillips, J. Leathwick, and J. Elith. 2016. Dismo: Species distribution modeling. R package ver. 1.0-15.

Hill, J. K., C. D. Thomas, and B. Huntley. 1999. Climate and habitat availability determine 20th century changes in a butterfly’s range margin. Proceedings of the Royal Society of London B: Biological Sciences 266:1197–1206.

Hines, J. E. 2006. Program presence. See http://www.mbrpwrc.usgs.gov/software/doc/presence/presence.html.

Hines, J. E., J. D. Nichols, and J. A. Collazo. 2014. Multiseason occupancy models for correlated replicate surveys. Methods in Ecology and Evolution 5:583–591.

Hines, J. E., J. D. Nichols, J. A. Royle, D. I. MacKenzie, A. Gopalaswamy, N. Kumar, and K. Karanth. 2010. Tigers on trails: Occupancy modeling for cluster sampling. Ecological Applications 20:1456–1466.

Johnson, D. S., P. B. Conn, M. B. Hooten, J. C. Ray, and B. A. Pond. 2013. Spatial occupancy models for large data sets. Ecology 94:801–808.

Kearney, M., and W. Porter. 2009. Mechanistic niche modelling: Combining physiological and spatial data to predict species ranges. Ecology Letters 12:334–350.

Kellner, K. 2015. jagsUI: A wrapper around rjags to streamline JAGS analyses. R package version 1.

Kéry, M. 2011. Towards the modelling of true species distributions. Journal of Biogeography 38:617–618.

Kéry, M., and J. A. Royle. 2015. Applied hierarchical modeling in ecology: Analysis of distribution, abundance and species richness in R and BUGS: Volume 1: Prelude and static models. Academic Press.

Kuo, L., and B. Mallick. 1998. Variable selection for regression models. The Indian Journal of Statistics, Series B 60:65–81.

Link, W. A., and J. R. Sauer. 2002. A hierarchical analysis of population change with application to Cerulean Warblers. Ecology 83:2832–2840.

Lunn, D. J., A. Thomas, N. Best, and D. Spiegelhalter. 2000. WinBUGS-a Bayesian modelling framework: Concepts, structure, and extensibility. Statistics and Computing 10:325–337.

MacKenzie, D. I., J. D. Nichols, J. E. Hines, M. G. Knutson, and A. B. Franklin. 2003. Estimating site occupancy, colonization, and local extinction when a species is detected imperfectly. Ecology 84:2200–2207.

MacKenzie, D. I., J. D. Nichols, J. A. Royle, K. H. Pollock, L. Bailey, and J. E. Hines. 2017. Occupancy estimation and modeling: Inferring patterns and dynamics of species occurrence. Elsevier.

Ntzoufras, I. 2002. Gibbs variable selection using BUGS. Journal of statistical software 7:1–19.

Pardieck, K. L., D. J. Ziolkowski Jr, M. Lutmerding, K. J. Campbell, and M.-A. R. Hudson. 2016. North american breeding bird survey dataset 1966-2015, version 2015.0. U.S. Geological Survey, Patuxent Wildlife Research Center doi:10.5066/F7W0944J.

Plummer, M. 2012. JAGS: Just another Gibbs sampler. Astrophysics Source Code Library.

R Core Team. 2016. R: A language and environment for statistical computing. R Foundation for Statistical Computing, Vienna, Austria.

Reich, B. J., J. S. Hodges, and V. Zadnik. 2006. Effects of residual smoothing on the posterior of the fixed effects in disease-mapping models. Biometrics 62:1197–1206.

Rich, J. L., and D. J. Currie. 2018. Are north american bird species’ geographic ranges mainly determined by climate? Global Ecology and Biogeography.

Rosenberg, K. V., T. Will, D. A. Buehler, S. B. Swarthout, W. E. Thogmartin, R. E. Bennett, and R. Chandler. 2016. Dynamic distributions and population declines of golden-winged warblers. Studies in Avian Biology 49:3–28.

Rushing, C. S., T. B. Ryder, A. L. Scarpignato, J. F. Saracco, and P. P. Marra. 2016. Using demographic attributes from long-term monitoring data to delineate natural population structure. Journal of applied ecology 53:491–500.

Sauer, J., J. Hines, J. Fallon, K. Pardieck, D. Ziolkowski, and W. Link. 2015. Breeding Bird Survey Summary and Analysis 1966-2013. Version 01.30.2015. USGS Patuxent Wildlife Research Center Laurel MD:http://www.mbr-pwrc.usgs.gov/bbs/bbs.html.

Tingley, M. W., and S. R. Beissinger. 2009. Detecting range shifts from historical species occurrences: New perspectives on old data. Trends in Ecology & Evolution 24:625–633.

Twedt, D. J. 2015. Estimating regional landbird populations from enhanced North American Breeding Bird Surveys. Journal of Field Ornithology 86:352–368.

Valpine, P.de, D. Turek, C. J. Paciorek, C. Anderson-Bergman, D. T. Lang, and R. Bodik. 2017. Programming with models: Writing statistical algorithms for general model structures with NIMBLE. Journal of Computational and Graphical Statistics 26:403–413.

White, G. C., and K. P. Burnham. 1999. Program MARK: Survival estimation from populations of marked animals. Bird Study 46:S120–139.

Wood, S. N. 2017. Generalized additive models: An introduction with R. CRC press.

